# Aberrant motor contagion of emotions in psychopathy and high-functioning autism

**DOI:** 10.1101/2021.07.05.450842

**Authors:** Lihua Sun, Lasse Lukkarinen, Tuomo Noppari, Sanaz Nazari-Farsani, Vesa Putkinen, Kerttu Seppälä, Matthew Hudson, Pekka Tani, Nina Lindberg, Henry K. Karlsson, Jussi Hirvonen, Marja Salomaa, Niina Venetjoki M., Hannu Lauerma, Jari Tiihonen, Lauri Nummenmaa

## Abstract

Psychopathy and autism are both associated with aberrant social interaction and communication, yet only psychopaths are markedly antisocial and violent. Here we compared the functional neural alterations underlying these two different phenotypes with distinct patterns of socioemotional difficulties. We studied 19 incarcerated male offenders with high psychopathic traits, 20 males with high-functioning autism and 19 age-matched healthy controls. All groups underwent functional magnetic resonance imaging while they viewed dynamic happy, angry and disgust facial expressions or listened to laughter and crying sounds. Psychopathy was associated with reduced somatomotor responses to almost all expressions, while subjects with autism demonstrated less marked and emotion-specific alterations in the somatomotor area. These data suggest that psychopathy and autism involve both common and distinct functional alterations in the brain networks involved in socioemotional processing.

## Introduction

Psychopathy is characterized by recurring antisocial behaviour, bold, disinhibited, and egotistical traits, and lacking empathy and remorse [1]. Psychopathy is causally linked with criminal behaviour and violence [2]. While the prevalence of psychopathy is around 1% in normal population, it is around 20% in incarcerated offenders [3] and 16.4% in Finnish incarcerated offenders [4]. Because the behavioural and emotional symptoms are persistent, psychopathy might have organic basis. Neuroimaging studies have found that psychopathic offenders have lower volume in the frontal cortex and in limbic regions including insula and amygdala [5–9]. These structural alterations are accompanied with abnormal responsiveness of the limbic system. Psychopathic subjects show weaker activity in amygdala and hippocampus, striatum and cingulate cortices while viewing emotional facial expressions. When viewing empathy-eliciting scenes, psychopathic individuals show significantly reduced frontocortical brain activity compared to healthy controls, consistent with their lowered care motivation [10, 11]. Conversely, stronger responses are observed in frontal cortical regions [12, 13], particularly when viewing violent emotional episodes [9]. The distorted limbic outputs combined with dysfunction in executive frontal cortical and social decision-making systems could thus predispose psychopaths to violent and antisocial behaviour.

Autism spectrum disorders (ASD) in turn are characterized by abnormal social interaction and communication, restricted interests, repetitive behaviour and sensory anomalies [14, 15]. ASD have variable clinical phenotypes from mild to severe, and even wider continuum of social-communicative ability extending into the general population has been proposed [16, 17]. Neuroimaging studies have linked ASD with aberrant structure and function in socioemotional brain networks such as those involved in processing of goal-directed actions and biological motion (superior temporal sulcus, STS), theory of mind (medial prefrontal cortex; mPFC and temporo-parietal junction; TPJ), and emotion (amygdala) [18, 19]. Similar to psychopathy, ASD is associated with reduced limbic activation by emotional stimuli such as happy, fearful and disgusted faces [20, 21], while a meta-analysis suggests that the aberrant emotional recognitions in ASD tend to be emotion-selective [18].

Although both psychopathy and autism are associated with abnormal socioemotional functioning and aberrant functioning in comparable brain systems have been implicated in both conditions, the behavioural phenotypes are markedly different. While psychopathy is consistently associated with severe antisocial and criminal behaviour, ASD is not systematically linked with increased likelihood for crime [22] despite aggression being somewhat common in autistic samples [23]. Furthermore, while psychopathic individuals can use their superficial charm and glib for manipulating other people [24], autistic individuals have in general severe difficulties in maintaining even routine social interactions. Comparison between autistic and psychopathic individuals provides a unique opportunity for addressing whether specific perturbations of the socioemotional brain networks are linked with distinct social and antisocial behavioural patterns. However, to our knowledge no prior study has directly compared functional brain basis of psychopathy and autism.

### The current study

In the current study we compared neural responses to emotional communicative signals in healthy controls versus incarcerated psychopaths and individuals with ASD. All groups underwent functional magnetic resonance imaging while they viewed dynamic happy, angry and disgust facial expressions or listened to laughter and crying sounds. We show that psychopathy is associated with reduced somatomotor responses to all expressions, while in ASD comparable alterations were found only for laughter and disgusted facial expressions.

Direct comparison revealed that downregulation of the somotomotor responses to all facial expressions was larger in psychopathy versus ASD. These data suggest that psychopathy and ASD involve both common and distinct functional alterations in the brain networks involved in socioemotional processing.

## Methods

### Subjects

We studied 19 convicted male offenders with high psychopathic traits, 20 males with high-functioning autism, and 19 age-matched healthy controls. The study was approved by the ethical committee of the Hospital District of Southwest Finland, and was conducted in accordance of the Helsinki declaration. All subjects completed informed consent forms prior to participating.

Offenders were inmates of the Turku Prison and they have been sentenced for murder (n = 5), manslaughter (n = 5), attempted manslaughter (n = 3) or grievous bodily harm (n = 6). Psychiatric diagnoses for offenders were based on prison health care, and forensic psychiatric violence risk assessments or most thorough forensic psychiatric examination reports concerning legal responsibility, two recruitment interviews and semi-structured PCL-R interviews. Final consensus diagnoses were made by two medical specialists (M.S., H.L.) both in psychiatry and forensic psychiatry with 13-25 years of experience in the field of prison psychiatry, which was also assisted by a psychologist (N.V.) with a 15-year working history in Psychiatric Hospital for Prisoners. Exclusion criteria were BMI more than 30, psychotic or other severe psychiatric illnesses, autoimmune illnesses, use of opioids, antipsychotic medication other than very small doses for insomnia, current substance abuse, exceptional risk of violence, claustrophobia, and other contraindications for functional imaging.

None of the offender group was psychotic nor suffered from a significant mood disorder. The group consisted of 16 subjects with antisocial personality disorders (ASPD) as defined by DSM-5 criteria [15], and three who did not fulfil the criterion of conduct disorder before age of 15 years but only the other criteria of antisocial personality. Of the later three, each happened to get the same result of 20/40 PCL-R points. One of them did not get any psychiatric diagnosis although posttraumatic stress disorder was suspected and he was the only subject of the inmate group who did not have any reported or documented history of substance abuse. Another of these three had abused alcohol, amphetamines and cannabis, and the third amphetamines, cannabis and steroids. Among the 16 subjects with diagnosis of ASPD, five were also diagnosed with Attention-Deficit/Hyperactivity disorder (ADHD), four with ICD-10 emotionally unstable personality, and one with panic disorder. Three subjects had history of poorly defined previous anxiety or depression. Among subjects with no diagnosed ASPD, there had been some previous signs of temporary adjustment disorders or insomnia in the prison with no clear diagnoses. History of excessive alcohol use was present in 13 subjects, and 18 subjects had self-reported or documented use of illegal substances including black market benzodiazepines, pregabalin or opioids, cannabis, amphetamines, gamma-hydroxybutyrate, MDPV, anabolic steroids and cocaine. Information concerning the severity of abuse were considered unreliable.

Psychopathy scores of the offenders were evaluated with semi-structured interview by experienced forensic psychiatrists or psychologists based on the Psychopathy Checklist-revised (PCL-R) [25]; psychopathy scores of healthy controls and autistic subjects were obtained via the Levenson Self-Report Psychopathy Scale (LSRP) questionnaire [26]. LSRP measures two dimensions of psychopathy with the primary psychopathy score indicating inclination to lie, lack of remorse, and callousness, and the secondary psychopathy score indicating impulsivity, short temper, and low toleration for frustration. Offenders were escorted by two prison guards to the local research institute for the brain imaging study. Clinical information of offenders is found in **supplementary Table S1**.

Subjects in the ASD group were volunteers from Helsinki and Turku University Hospital Neuropsychiatric Clinic; one subject was also recruited from Neuropsychiatric Clinic Proneuron, in Espoo. The ASD diagnoses were verified by research psychologist, neurologist and psychiatrist following DSM-5 criteria based on patient history, all accessible information from births records, well-baby clinics and school healthcare. An additional current Autism Diagnostic Observation Schedule (ADOS) assessment [27] was also used to clarify ASD diagnostics. All ASD subjects were diagnosed with ASD, with six also diagnosed with ADHD and eight with other mood and anxiety disorders. Healthy subjects and ASD group also completed the Autism-Spectrum Quotient (AQ) questionnaire [28]. None of the ASD subjects had current severe mental disorder, as assessed by SCID-I interview [29]. Dopaminergic medications (antipsychotics, psychostimulants, bupropion) were withdrawn before measurements but those who had SSRI-medication could not be withdrawn. Clinical information of the ASD subjects is found in **supplementary Table S2**.

### Facial expression task

In the emotional facial expression task (**Fig 1A**), subjects viewed short video clips (5 s) of dynamic facial expressions of joy, disgust and anger selected from ADFES database [30]. All clips begun with a neutral face, followed by a dynamic display of the facial expression. Prior to each clip, subjects were shown the first frame of the video (i.e. neutral face) for 3.5 s to avoid peaks in low-level visual activation due to simultaneous visual stimulus and motion onset. This was followed by the dynamic expression from neutral to full expression, with the full-blown phase held until the end of the clip. Each stimulus was followed by a random 4 - 8 s of rest period. Again, to avoid peaks in low-level visual cortical activations, a scrambled picture of the upcoming model was shown during the rest period. To keep subjects focused on the task, four trials (out of 36 trials in total) contained a still picture of the neutral face instead of the video clip. Subjects were asked to press the response button as soon as they detected a trial without any facial motion. These trials were excluded from the analysis.

**Figure 1.**
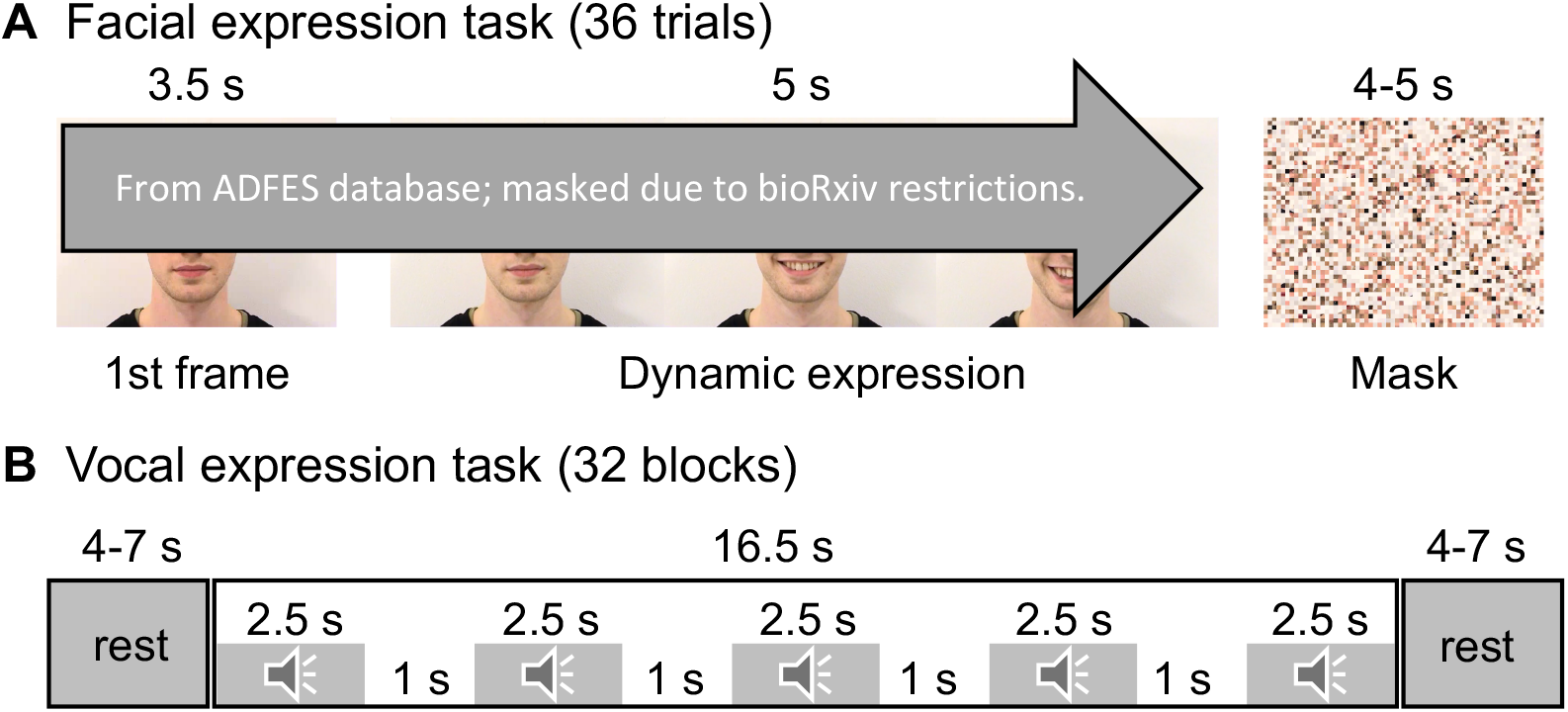
Experimental design for the facial expression task (A) and vocal expression task (B).

### Vocal expression task

In the vocal expression task (**Fig 1B**), the subjects listened to short laughter and crying vocalizations and control stimuli that were generated by time-domain scrambling of the original sounds. The original stimuli have been validated and described in detail in [31]. The experiment was run using a blocked design. In each 16.5s block, five 2.5s stimuli from one category (i.e., laughter, crying sounds, scrambled laughter, or scrambled crying sounds) were played with a 1s silent period between stimuli. Order of the blocks were randomized. The blocks were interspersed with rest blocks lasting for 4-7s. To keep participants focused on the task, an animal sound (sound of vocalization of an alpaca) was presented randomly during 50% of the rest blocks. The subjects were instructed to press the response button whenever they heard the alpaca, and the behavioural outcomes were inspected for the focus of attention. A total of 32 blocks (8 blocks per stimulus type) were run.

### fMRI acquisition and preprocessing

The MRI data were acquired with Phillips Ingenuity TF PET/MR 3T whole-body scanner. High-resolution (1 mm^3^) structural brain images were acquired using a T1-weighted sequence (TR 9.8 ms, TE 4.6 ms, flip angle 7°, 250 mm FOV, 256 × 256 reconstruction matrix). Radiologist screened the images for structural abnormalities. Functional data were acquired using a T2*-weighted echo-planar imaging sequence (TR = 2600 ms, TE = 30 ms, 75° flip angle, 240 mm FOV, 80 × 80 reconstruction matrix, 62.5 kHz bandwidth, 3.0 mm slice thickness, 45 interleaved slices acquired in ascending order without gaps). A total of 206 (facial expression task) or 290 (laughter task) functional volumes were acquired. We used fMRIPrep 1.3.0.2 to preprocess MRI data [32]. Anatomical T1-weighted (T1w) reference images were processed following steps: correction for intensity non-uniformity, skull-stripping, brain surface reconstruction, spatial normalization to the ICBM 152 Nonlinear Asymmetrical template version 2009c [33] using nonlinear registration with antsRegistration (ANTs 2.2.0) and brain tissue segmentation. Functional MRI data were processed following steps: coregistration to the T1 reference image, slice-time correction, spatial smoothing with an 6mm Gaussian kernel, automatic removal of motion artifacts using ICA-AROMA [34] and resampling to the MNI152NLin2009cAsym standard space. Quality of images was assessed via the visual reports of fMRIPrep and inspected manually in accord to whole-brain field of view coverage, proper alignment to the anatomical images, and signal artifacts. All functional data were restrained in the analysis.

### Full-volume GLM data analysis

The fMRI data were analysed in SPM12 (Wellcome Trust Center for Imaging, London, UK, (http://www.fil.ion.ucl.ac.uk/spm). The whole-brain random effects model was applied using a two-stage process with separate first and second levels. For each subject, GLM was used to predict regional effects of task parameters on BOLD indices of activation. In the facial expression task, contrast images were generated for dynamic happy, angry, or disgusted facial expressions versus static neutral faces (i.e., the initial 3.5s of each video without motion) and subjected to second-level analyses for population level inference. In the vocal expression task, contrast images were generated for laughter or crying sound versus corresponding scrambled sounds and subjected to second-level analyses. We first tested the task-dependent activations in each group and conducted the between-groups comparisons for each effect of interest. Statistical threshold was set at p < 0.05, FDR corrected at cluster level.

### Region of interest (ROI) analysis

To visualize the between-groups differences, BOLD signals in anatomically defined regions of interest (ROIs) were also analysed. ROIs were selected considering their important roles in emotional processing and also in accord to findings of the full volume analysis. These ROIs included anterior, middle and posterior cingulate cortex, and precuneus defined by the AAL atlas [35]. We also included the subregions of motor area parcelled in the Juelich Atlas with masks generated using the SPM Anatomy toolbox [36]. These subregions include the primary motor cortex (M1) corresponding to Brodmann areas (BA) 4a and 4b, the supplementary motor area (M2) corresponding to BA6 [37] the primary somatosensory cortex (S1) including BA3a, BA3b, BA1 and BA2 [38, 39], and the secondary somatosensory cortex (S2) including parietal operculum 1-4 [40]. Regional beta weights were estimated from first-level contrast images of each subject using the MarsBaR toolbox [41]. ROI data were analysed using two-sample t-test using with R statistical software (version 3.6.3).

## Results

### Psychopathy and autism evaluation in the studied groups

Basic subject information is summarized in **Table 1**. There was no statistical difference between ASD group and controls for LSRP primary psychopathy scores (t = 1.20; p = 0.24), but ASD group had slightly higher secondary psychopathy score (t = 3.12; p = 0.003). ASD group had higher AQ score compared to controls (t = 11.14; p < 0.001)).

**Table 1.**
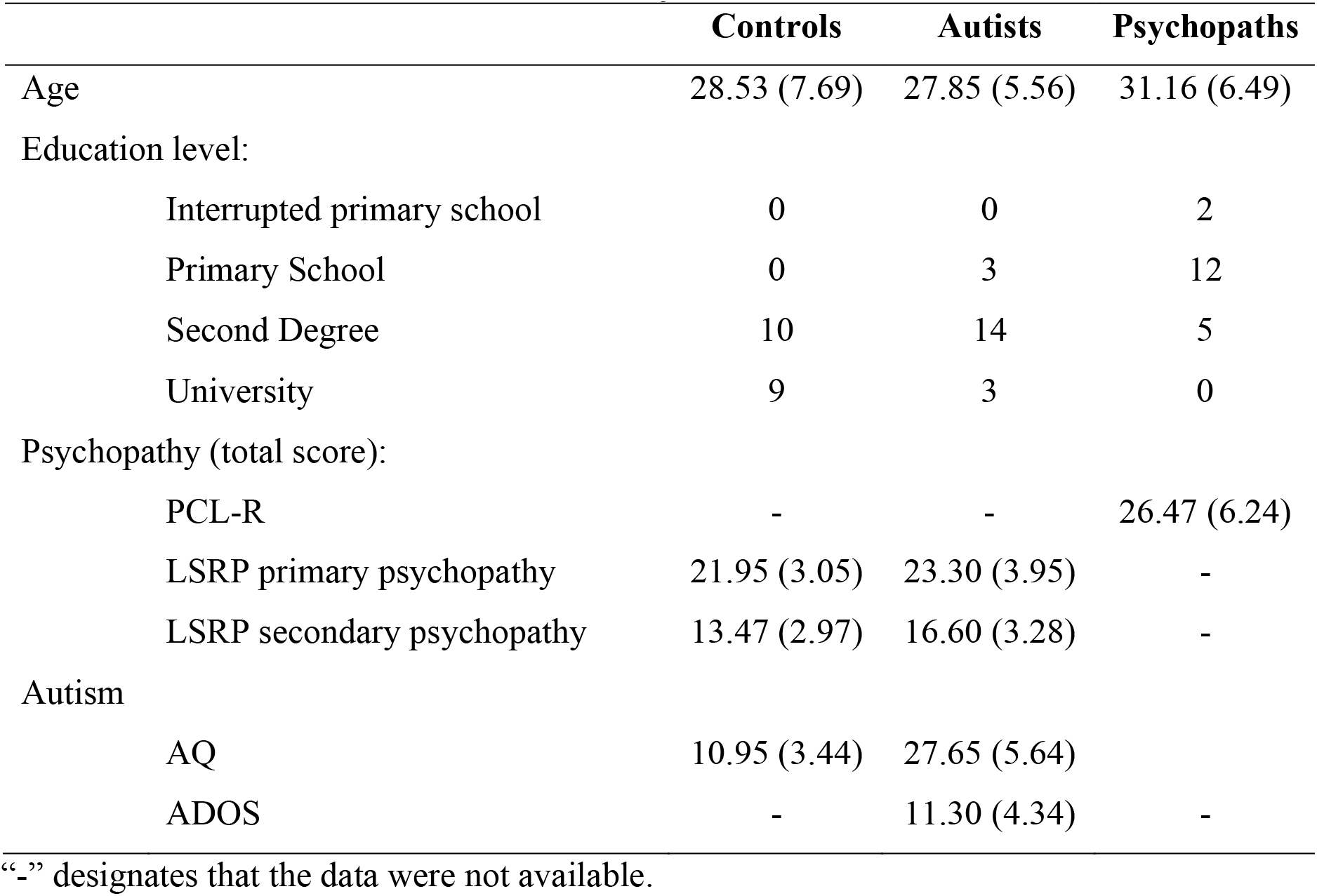
Basic subject characteristics.

### Regional responses to positive emotional stimuli

In control subjects, happy faces elicited activation in occipital cortex, fusiform gyrus (FG), cingulate cortex (CC), motor area including the primary (S1) and secondary (S2) somatosensory cortex and primary (M1) and supplementary motor (M2) areas, medial frontal cortex (MFC), middle (MTG) and superior temporal gyrus (STG), precuneus, cuneus, amygdala, hippocampus, striatum and thalamus (**Fig 2A**). Social laughter sounds elicited activation in primary and secondary auditory cortices, CC, motor area, MFC, MTG and STG, precuneus, amygdala, hippocampus, striatum and thalamus. These activations by both happy faces and laughter were weakened in the autistic individuals and markedly abolished in the psychopathic individuals, with the exception of the temporal activations (**Fig 2A**). In psychopaths, large-scale deactivation was also observed for laughter.

**Figure 2.**
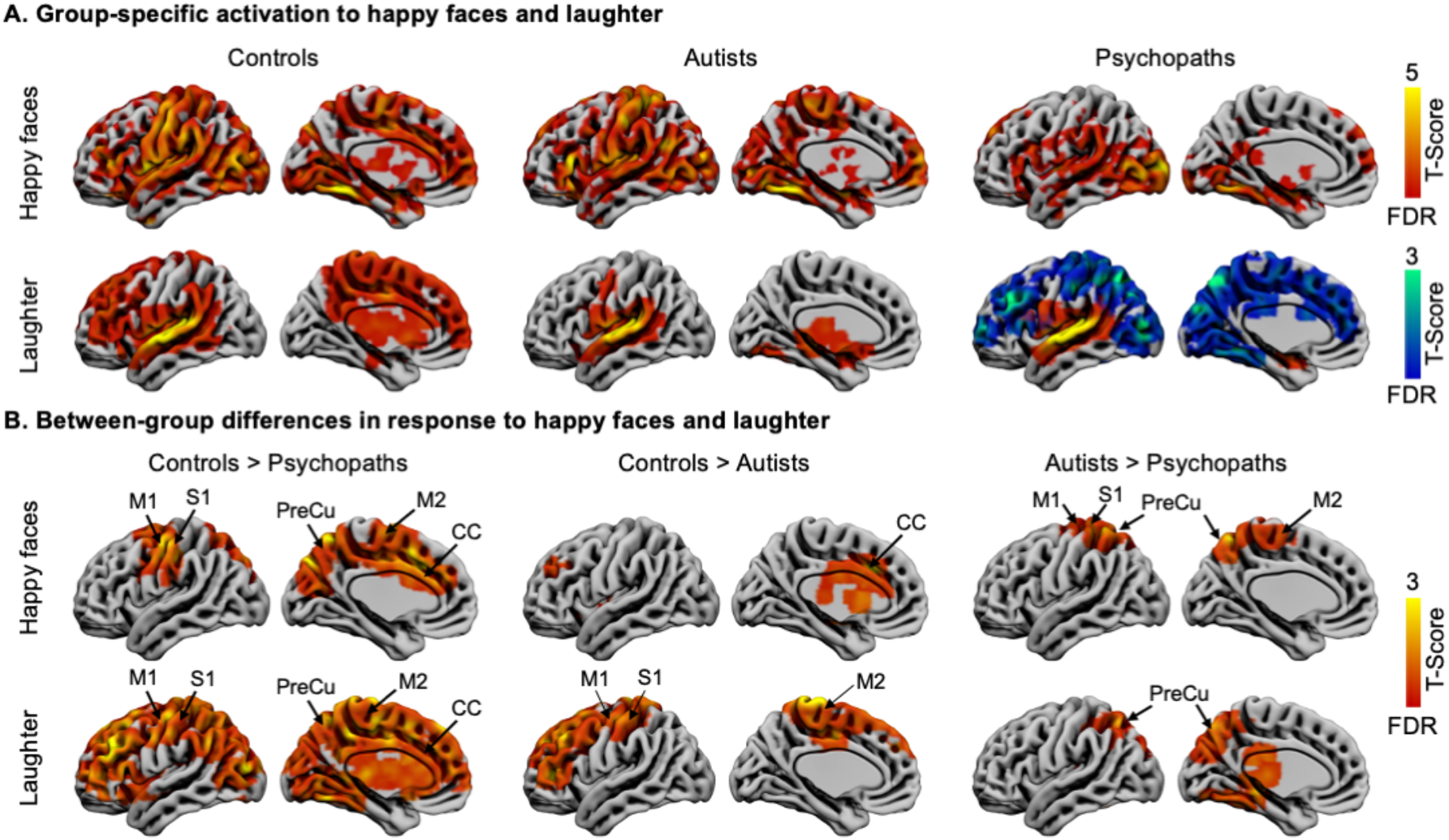
Brain responses to happy faces and social laughter. **A.** Responses to happy faces and laughter separately for each group. Hot colour indicates activation and cool colour indicates deactivation. **B.** Between-group differences in response to happy faces and laughter. Data are thresholded at p < 0.05 with FDR cluster-level correction. CC = cingulate cortex, S1 = primary somatosensory cortex, S2 = secondary somatosensory cortex, M1 = primary motor cortex, M2 = supplementary motor area, PreCu = precuneus; left hemispheres were presented for visualization.

This was confirmed in direct between-group contrasts (**Fig 2B**). Compared to controls, psychopathic subjects showed dampened responses to happy faces and laughter in motor area, CC and precuneus. Dampened activation in response to laughter expanded largely to frontal and posterior brain areas and subcortical regions. Compared to controls, autists showed dampened responses in the middle and anterior CC to happy faces, and in the motor area (also expanding frontally) to laughter. Compared to autists, psychopaths showed dampened response in motor area to happy faces, and in precuneus to laughter.

### Regional responses to negative emotional stimuli

In controls, both angry and disgusted faces elicited activation in the occipital cortex, FFA, CC, the motor area, MFC, MTG and STG, precuneus, amygdala, hippocampus, striatum, and thalamus (**Fig 3A**). Comparable activation of these regions was found in autistic individuals, while similarly as for happy faces and laughter, activity in these brain regions in psychopathic individuals were markedly abolished (**Fig 3A**).

**Figure 3.**
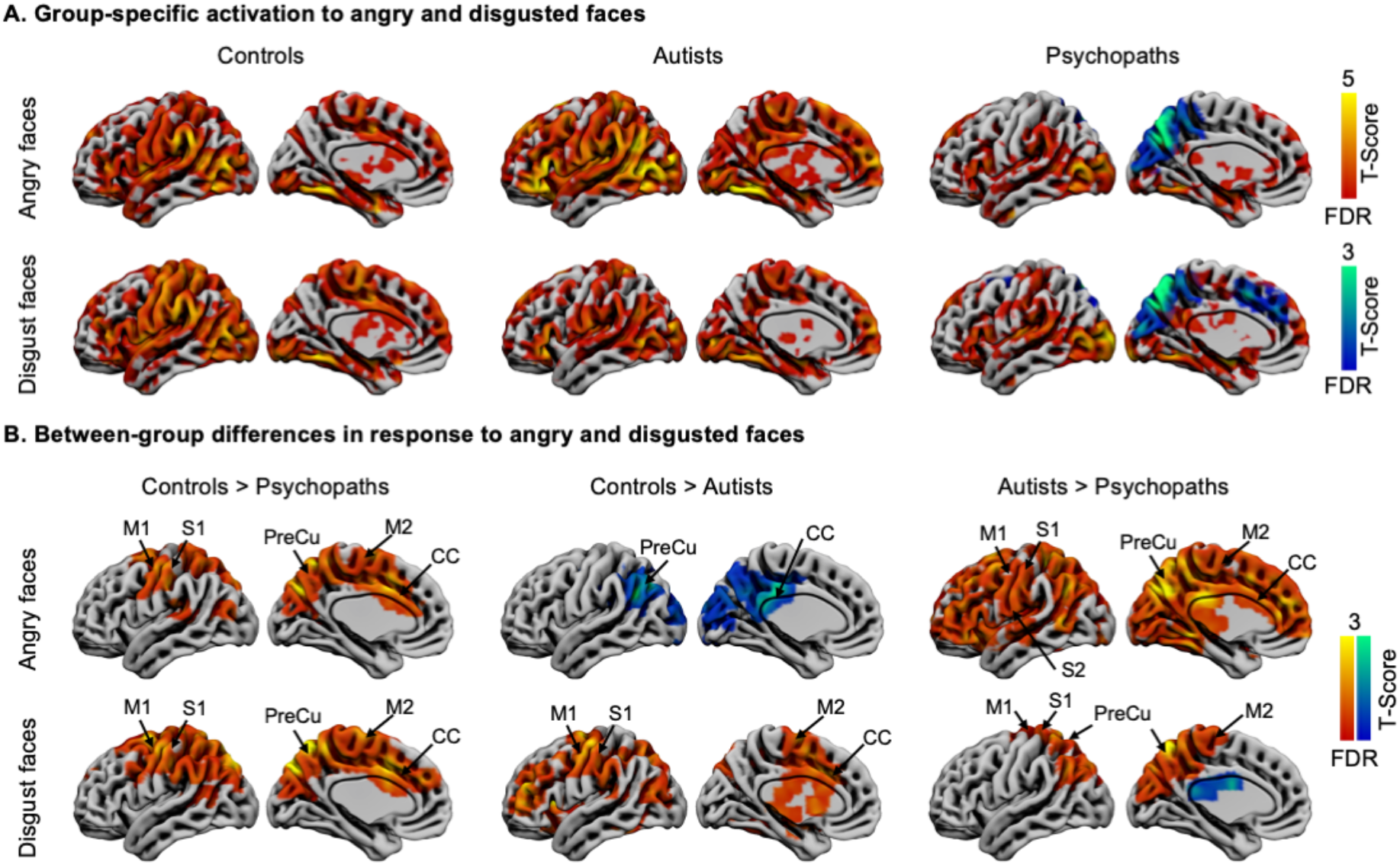
Brain responses to angry and disgusted faces. **A.** Responses to angry and disgusted faces separately for each group. **B.** Between-group differences in responses to angry and disgusted faces. Data are thresholded at p < 0.05 with FDR cluster-level correction. CC = cingulate cortex, S1 = primary somatosensory cortex, S2 = secondary somatosensory cortex, M1 = primary motor cortex, M2 = supplementary motor area, PreCu = precuneus; left hemispheres were presented for visualization.

This was also confirmed in direct between-groups contrast (**Fig 3B**). Compared to controls, psychopaths showed dampened activation in CC, motor area and precuneus to both angry and disgusted faces. Compared to controls, autists also showed dampened activation in the CC, motor area and precuneus to disgusted faces. However, in response to angry faces, autists showed increased activation in precuneus and posterior CC. Compared to autists, psychopaths showed global deactivation to angry faces, and damped activation in motor area and precuneus to disgusted faces.

Crying sound elicited activation mainly in the primary and secondary auditory cortices, and nearby regions (**supplementary Fig S1**). However, group comparisons did not show statistical differences.

### Region of interest analysis

Region of interest (ROI) analysis demonstrated between-group differences that were in accord with the full-volume analysis (**Fig 4**). In response to laughter (**Fig 4A**), psychopaths showed reduced activation in M1, M2, S1, and the whole motor area (combined of M1, M2, S1 and S2), anterior (ACC) and middle cingulate cortex (MCC), compared to controls. There were no statistically significant differences between controls and autists, although numerically mean activity was strongest in controls and weakest in psychopaths in most ROIs.

**Figure 4.**
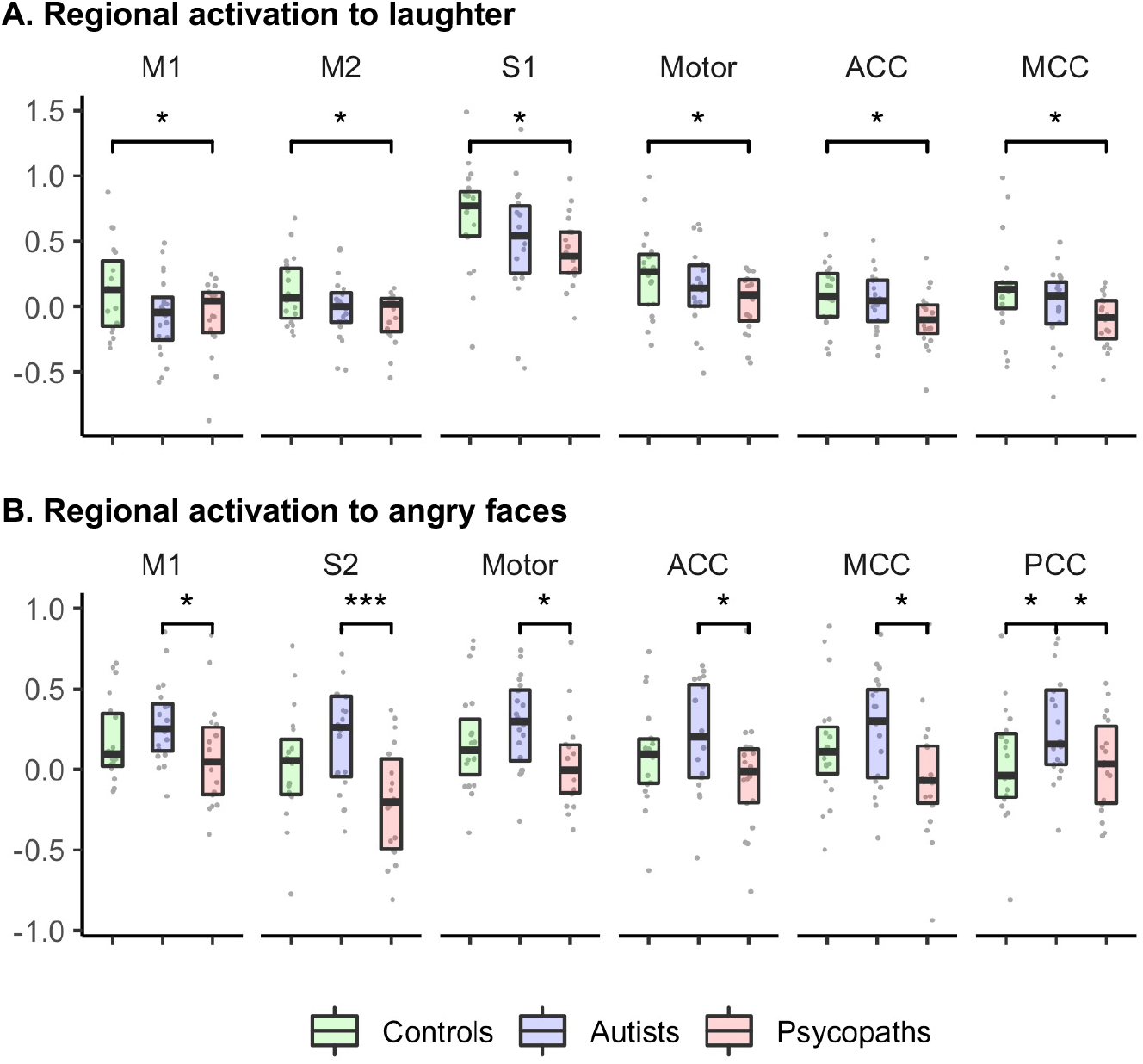
Region-of interest analysis for laughter and angry faces. Between group comparisons were conducted using student’s t test, with significance levels marked: *p < 0.05, ***p < 0.01. M1 = primary motor cortex, M2 = supplementary motor area, S1 = primary somatosensory cortex, S2 = secondary somatosensory cortex, Motor = combined region of M1, M2, S1 and S2, ACC = anterior cingulate cortex, MCC = middle cingulate cortex, PCC = posterior cingulate cortex; data involved both hemispheres.

In response to angry faces (**Fig 4B**), psychopaths demonstrated reduced activation in ROIs including the M1, S2, whole motor area, ACC, MCC, and PCC compared to autists. Autists also showed increased activation in the PCC compared to controls. No between-group differences were found for crying sounds and happy faces. To disgust faces, psychopaths showed reduced activation in MCC compared to controls (data not shown).

## Discussion

Incarcerated psychopaths and high-functioning autistic subjects showed both common and unique alterations in the brain responses to positive and negative facial and vocal social communicative signals. Compared with controls, psychopaths showed supressed brain activation toward all communicative signals except crying sounds. Weaker activity was observed in somatosensory, motor and cingulate cortex. This effect was less pronounced in the autistic subjects and was observed primarily for laughter and disgusted facial expressions. Direct comparison between psychopaths and subjects with ASD revealed that the somatomotor responses were weaker in psychopaths. Altogether, our data show that alterations in somatomotor processing of emotional signals is a common characteristic of criminal psychopathy and autism, yet the degree and specificity of these alterations distinguishes between these two groups.

Our main finding was that somatomotor “mirroring” of vocal and facial emotional expressions was altered in both criminal psychopaths and autists, and the somatosensory and motor responses to emotional signals were more reduced in the criminal psychopaths than in autistic subjects. This accords with previous studies that have found reduced brain activation during passive observation of others’ distress [11] or affective memory tasks [13] in psychopaths. Psychopathic offenders also show less behavioural contagion of laughing and yawning [42], and recent structural imaging study demonstrated that both criminal psychopathy as well as psychopathy-like traits in healthy controls are associate with lower volume in the somatosensory cortices [9].

Seeing others in a particular emotional state often triggers automatically the corresponding behavioural and somatic representation of that emotional state in the observer [43, 44]. Neuroimaging studies have confirmed that such somatomotor contagion of emotions is subserved by common neural activation for perception and experience of states such as pain [45–47], disgust [48] and pleasure [49], allowing to ‘tune in’ or ‘sync’ with other individuals [50, 51]. Furthermore, damage to somatosensory cortex [52], and their inactivation by transcranial magnetic stimulation [53] also impairs recognition of emotions from facial expressions. The widespread aberrant responsivity of the somatosensory cortex in psychopaths may explicate their asocial character, lack of empathy and egotistical traits [1]. As mirroring of others’ emotions and particularly distress plays a crucial role in empathy and inhibition of violent behaviour [54, 55], impaired somatic and motor contagion of others’ emotions may render psychopaths susceptible to antisocial behaviour and violence.

The autistic subjects also had lowered somatomotor responses to emotional signals, although this effect was less profound than in the psychopaths. Prior work shows that autistic individuals have difficulties in recognizing specific emotions [18, 19, 56], as well as difficulties in automatic mimicking facial expressions [57, 58]. Some studies have also shown that autistic subjects have deficient motor intention understanding ability, possibly linked with aberrant motor cognition [59–61]. In line with these studies, functional neuroimaging experiments show that brains of high-functioning autists do not synchronize with those of others while viewing naturalistic social interaction, indicative of aberrant automatic tuning in with others’ mental states [62, 63]. The present data highlight how the aberrant activity of the somatosensory and motor cortices may also contribute to these impairments.

Although both psychopathic and autistic subjects showed, in general, reduced responses to the vocal and facial emotional expressions, the specific patterns of these alterations differed across the groups. Overall, the emotional expressions evoked weaker responses in the psychopathic than autistic individuals, and for all facial expressions this effect was observed in the primary somatosensory, primary and supplementary motor cortex. Because motor responses to social communicative signals are fundamental for establishing social bonds between individuals [64, 65] and important for the formation of empathic responses [66–68], the present observed aberrant motor contagion may reflect the shared component of the socioemotional deficits in autism and psychopathy. Laughter expressions elicited to large-scale deactivation outside the auditory cortices only in psychopaths. Laugher is a universally recognized prosocial signal that is uses for bonding purposes, rather than an expression of positive emotional state [69], and many of the characteristics defining psychopathy are related to abnormal socioemotional interaction. It is thus possible that the aberrant neural responses to bonding signals such as laughter could link with the antisocial traits in psychopathy. Additionally, for angry faces, the difference between psychopathic and autistic subjects was markedly widespread, with psychopaths showing significantly reduced responses across the medial and lateral frontal cortices in comparison with the autistic subjects. These data suggest that autism-associated hypersensitivity of the neural systems responding to anger and hyposensitivity to prosocial cues such as laugher may explain the distinct patterns of social interaction and communication deficits in psychopathy and autism.

The current study also bears limitations. Although we aimed at recruiting prisoner subjects not using antipsychotics, antidepressants or anxiolytics, it was not possible to recruit a completely drug-naïve sample. The convicted offenders and healthy controls and autistic subjects also differ from each other regarding the available quality and quantity of social interaction, leisure time activities, education levels and so forth. Ideally, this kind of study should thus also involve a forensic but non-psychopathic sample. Our data are cross-sectional in nature and cannot resolve the potential causal link between the functional alterations and psychopathy and autism. Also, we only included male subjects in the current study and findings may not generalize to females.

In summary, our findings suggest that aberrant neural activity in somatomotor areas may be a common mechanism underlying the asocial behaviour in psychopathy and autism, while its severity and selectivity in response to different types of social communicative signals set these disorders apart. These data suggest that distinct conditions associated with social information processing abnormalities might share common neurobiological substrates despite distinct behavioural and clinical phenotypes.

## Acknowledgment

The study was supported by the Academy of Finland (grants no. 294897 and 332225), and European Research Council starting (grant 313000 to L.N.), Valon Vuoksi Foundation (grants to L.S. and L.L.). L.S. is personally supported by the Turku Collegium for Science and Medicine, University of Turku. We thank director Juhani Järvi and other staff members of Turku Prison who made it possible to safely guard and transport the inmates during the neuroimaging.

## Conflict of interest

The authors declare no conflict of interest.

## Notes

### Competing Interest Statement

The authors have declared no competing interest.

