## Supplementary data for "Aberrant motor contagion of emotions in psychopathy and high-functioning autism"

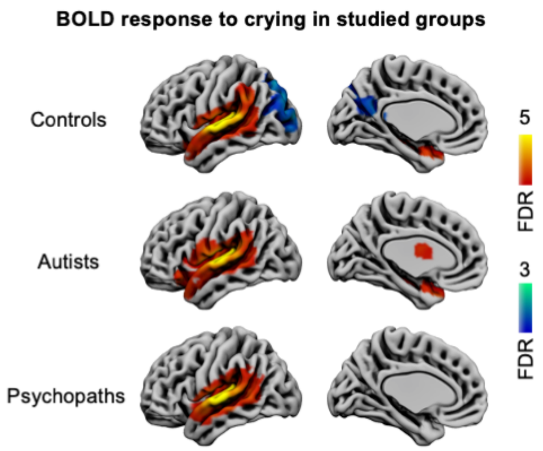


**Figure S1.** Brain responses to crying sound in the studied groups. There were no between-group differences. Results are thresholded at $p < 0.05$ with FDR cluster-level correction.

**Table S1.** Clinical characteristics of the violent offenders.

| Subject | Medication | Times of imprisonment | PCL-R Score |
|---|---|---|---|
| 1 | Hydroxyzine, Fluoxetine, Quetiapine* | 1 | 28 |
| 2 | Levothyroxine, Amitriptyline | 1 | 29 |
| 3 | Citalopram, Buspirone, Propranolol | 1 | 36 |
| 4 | Hydroxyzine, Melatonin | 2 | 28 |
| 5 | Melatonin, Hydroxyzine, Escitalopram*, Quetiapine*, Levomepromazine* | 1 | 29 |
| 6 | None | 1 | 20 |
| 7 | None | 4 | 27 |
| 8 | Hydroxyzine | 4 | 33 |
| 9 | Hydroxyzine | 6 | 33 |
| 10 | Mirtazapine | 1 | 21 |
| 11 | Melatonin, Buspirone*, Quetiapine* | 2 | 23 |
| 12 | Quetiapine, Propranolol, Amitriptyline, Hydroxyzine | 1 | |
| 13 | Quetiapine, Amitriptyline | 2 | 20 |
| 14 | None | 1 | 20 |
| 15 | Hydroxyzine, Buspirone, Risperidone, Atomoxetine, Atomoxetine | 1 | |
| 16 | Melatonin, Hydroxyzine, Propranolol, Quetiapine* | 3 | 30 |
| 17 | Melatonin | 4 | 24 |
| 18 | Hydroxyzine, Cetirizine, Melatonin, Amitriptyline, Mirtazapine, Venlafaxine | 1 | 16 |
| 19 | None | 9 | 35 |

*Stopped less than 14 days before brain imaging.

**Table S2.** Clinical characteristics of the ASD subjects.

| Subject | Medication | LSRP Primary Score | LSRP Secondary Score | AQ Score | ADOS Score |
|---|---|---|---|---|---|
| 1 | None | 24 | 18 | 37 | 16 |
| 2 | None | 26 | 16 | 27 | 7 |
| 3 | None | 18 | 14 | 31 | 10 |
| 4 | None | 29 | 14 | 30 | 18 |
| 5 | Melatonin | 19 | 15 | 35 | 7 |
| 6 | Fluoxetine | 29 | 15 | 25 | 15 |
| 7 | None | 22 | 19 | 28 | 7 |
| 8 | Zolpidem* | 30 | 14 | 29 | 16 |
| 9 | None | 31 | 21 | 27 | 9 |
| 10 | Levothyroxine, Cetirizine, Escitalopram | 22 | 14 | 32 | 7 |
| 11 | Melatonin | 22 | 16 | 19 | 12 |
| 12 | None | 25 | 23 | 32 | 13 |
| 13 | None | 20 | 19 | 27 | 14 |
| 14 | Venlafaxine | 24 | 21 | 22 | 2 |
| 15 | None | 20 | 12 | 33 | 12 |
| 16 | None | 22 | 10 | 18 | 6 |
| 17 | Escitalopram | 19 | 17 | 26 | 17 |
| 18 | Vortioxetine, Bupropion* | 20 | 18 | 24 | 14 |
| 19 | Melatonin | 24 | 20 | 34 | 14 |
| 20 | None | 20 | 16 | 17 | 14 |

*Stopped less than 14 days before brain imaging.
